# A single-cell view of human tissue aging reveals architectural decline beyond cellular composition

**DOI:** 10.64898/2026.09.06.749715

**Authors:** Ernesto Abila, Yimin Zheng, Zsuzsanna Bago-Horvath, André F. Rendeiro

**Affiliations:** CeMM Research Center for Molecular Medicine of the Austrian Academy of Sciences, Lazarettgasse 14 AKH BT 25.3, 1090, Vienna, Austria; Ludwig Boltzmann Institute for Network Medicine at the University of Vienna, Augasse 2-6, A-1090 Vienna, Austria; Department of Pathology, Medical University of Vienna, Währinger Gürtel 18-20, 1090 Vienna, Austria

**Keywords:** aging, histopathology, deep learning, single-cells, tissue biology

## Abstract

Aging reshapes the human body at the cellular level, yet how cell identity, morphology, and spatial organization remodel across organs and the adult lifespan remains poorly mapped, at a scale and lifespan coverage that molecular spatial assays cannot yet reach. Treating the GTEx histopathology archive as a population-scale, life-span-resolved resource for spatial biology, we detected over 3.5 billion single cells across 16 human organs from nearly one thousand individuals. Cell density declined pervasively but organ-specifically, and vision-language phenotyping resolved epithelial cells into nine subtypes with divergent aging trajectories, including loss of ovarian granulosa cells at ∼45% per decade. Community detection on spatial cell graphs identified functional tissue units, over a quarter of which remodeled with age along a shared trajectory from dense, specialized units toward sparser, stromal- and immune-enriched structures. Critically, this architectural remodeling was largely decoupled from cell composition (R^2^=0.07), showing that human tissues age along two partly independent axes, a pervasive loss of cells and a distinct remodeling of the architecture they form, with structural decline exceeding what cellular composition alone predicts.

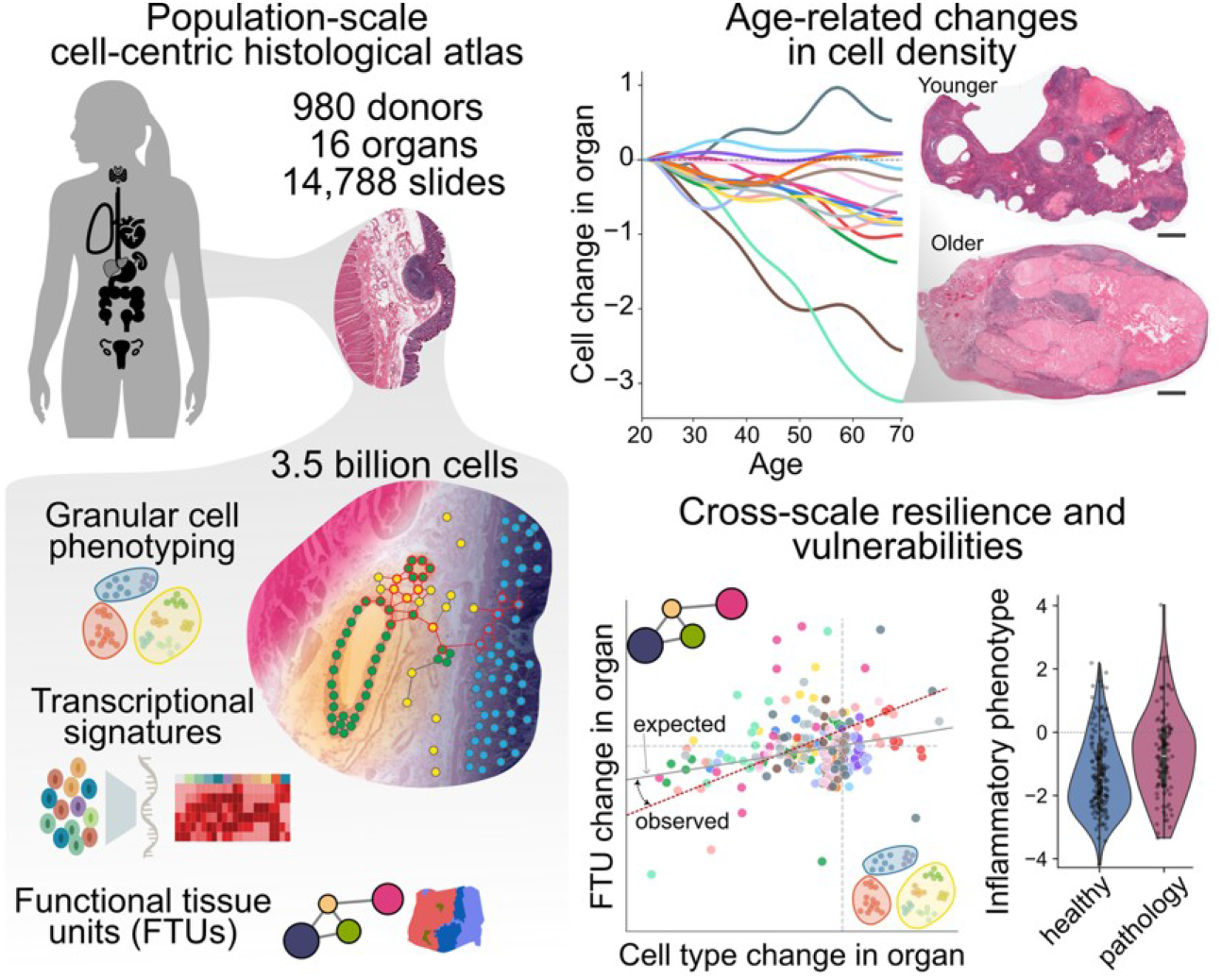

## Introduction

The physiology of the human body emerges from the coordinated arrangement of cells into functional tissue units^1^ such as intestinal crypts, ovarian follicles, and pulmonary alveoli, whose structural integrity is essential for organ function^2–4^. Cells occupy defined positions within this microanatomical context, and perturbations of the cellular microenvironment, from extracellular matrix remodeling to altered cell-cell interactions and disruption of cellular morphology, tissue architecture, and ultimately organ physiology^5,6^. Such architectural dysregulation underlies diseases ranging from cancer to chronic inflammatory conditions such as gastritis and esophagitis^7^.

Aging is a prominent example of progressive architectural remodeling that compromises organ function. Molecular hallmarks accumulating over the lifespan, including genomic instability, epigenetic alterations, and chronic inflammation, eventually manifest in the cellular morphology and tissue organization that underlie functional decline and age-related disease^8–10^. As populations age and the burden of these diseases grows worldwide^11,12^, resolving how cell composition, morphology, and tissue organization shift across human organs throughout the lifespan has become a pressing priority^10,13,14^.

Large-scale molecular atlases such as the Human Cell Atlas^14^ and HuBMAP^15^ have transformed our understanding of cellular diversity and its spatial organization. Yet the scale of tissue sampling, coverage of the adult lifespan, and systematic multi-organ comparison accessible to these technologies remain limited, leaving the population-scale, lifespan-resolved architecture of human tissues largely uncharted. Hematoxylin and eosin (H&E) histopathology, the standard assay in pathology, instead captures tissue morphology and architecture at single-cell resolution across entire cross-sections, and advances in computational pathology now make it interrogable at scale, from nuclear instance segmentation of individual cells^16–21^ to transferable, interpretable representations from foundation and vision-language models such as PLIP^22^ and CONCH^23^.

Building on these capabilities, we previously showed that population-scale histopathology archives can be activated to quantify tissue-level aging signatures^24^ and that recurrent microanatomical domains undergo selective remodeling across the lifespan^25^. What remains missing is a cellular-resolution view bridging these scales, linking the morphology and identity of individual cells to the functional tissue units they compose, to resolve how the cellular-to-architectural axis is reshaped during aging.

## Results

### A billion-scale cellular census reveals widespread loss of cellular density with age

To systematically characterize cellular and architectural aging, we leveraged whole-slide histopathology images (WSIs) from the Genotype-Tissue Expression (GTEx) cohort, spanning 16 organs from 980 donors aged 20 to 70 (14,788 WSIs; **Fig. 1a, Fig. S1a, b**). We segmented and broadly classified approximately 3.5 billion cells (epithelial, neoplastic, immune, connective) with CellViT^19^. Because cancer-trained models alone cannot resolve the cellular diversity of healthy tissues, we used this output as the entry point for a multi-modal phenotyping strategy combining cell morphology (CellViT^19^), nuclear morphology (DINOv2^26^), and vision-language embeddings of the surrounding tissue context (PLIP^22^), enabling both finer phenotyping and orthogonal quality control of the initial labels (**Fig. 1a**).

**Figure 1:**
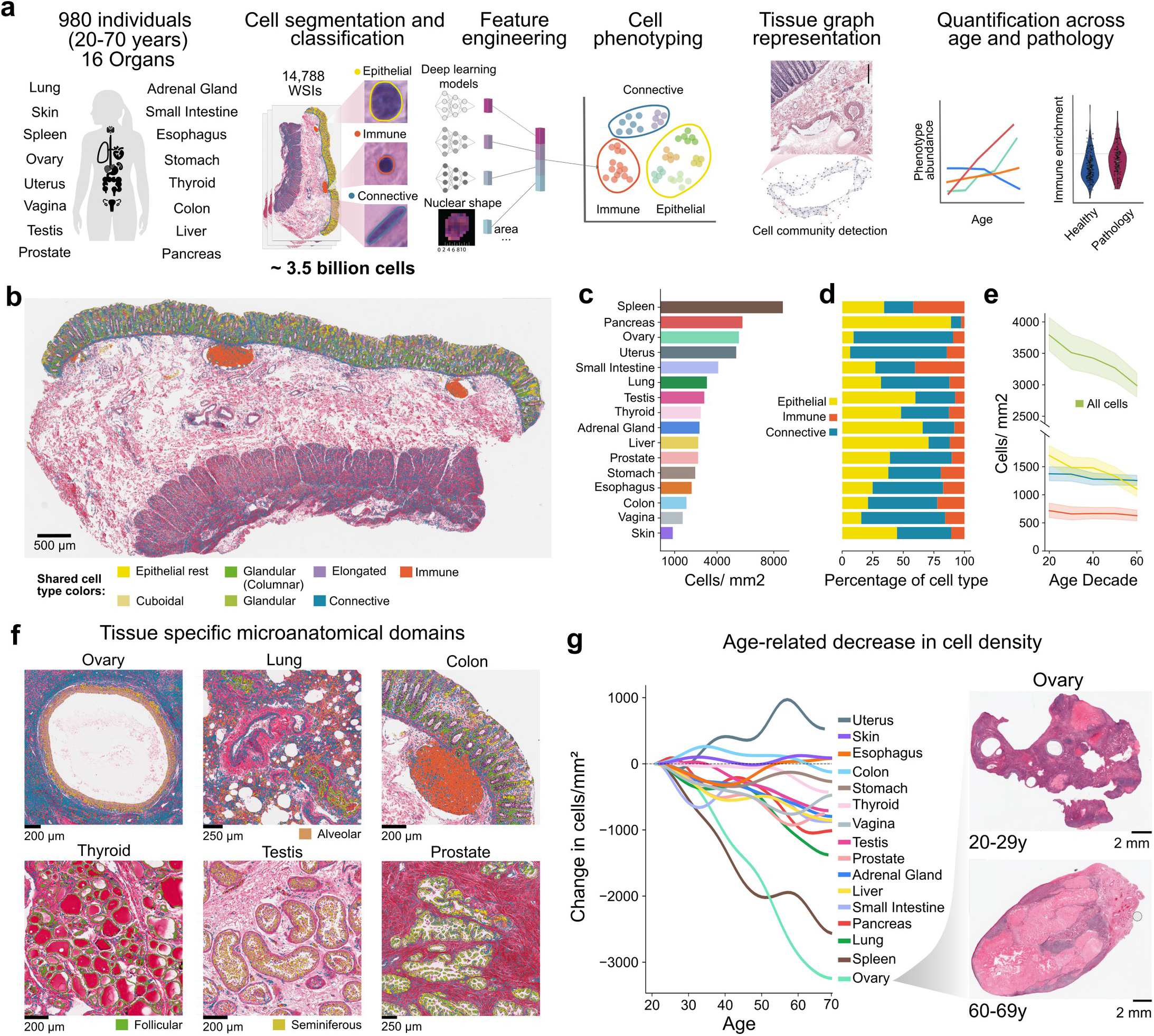
Single-cell atlas of 16 human organs reveals age-associated changes in cell composition and tissue architecture. **a)** Study design: 16 organs from 980 donors (age 20-70; 14,788 WSIs), with segmentation and classification of ∼3.5 billion cells into epithelial, immune, and connective types, followed by multi-modal feature engineering (CellViT, DINOv2, CONCH), phenotyping, spatial community detection, and quantification of age-associated and pathological change. **b)** Example colon WSI with cells colored by type. **c)** Mean cell density (cells/mm2) per organ. **d)** Cell type composition per organ (stacked proportions of epithelial, immune, and connective cells). **e)** Cell density distributions across age decades per cell type. **f)** Representative crops from six organs with cells colored by type (as in b). **g)** Smoothed cell density across age per organ, with representative young and old donor ovary WSIs.

An example colon section (**Fig. 1b**) illustrates that these classifications resolve the canonical layers of the intestinal wall, from the crypt-bearing mucosa and its lamina propria immune cells, through submucosal lymphoid aggregates and vascular networks, to the smooth muscle of the muscularis propria. Across organs, cell density varied more than eightfold, from the spleen as the densest (>8,000 cells/mm ^2^) to the skin as the sparsest (<1,000 cells/mm^2^) (**Fig. 1c**). Cell type composition likewise tracked physiological role, with the pancreas most epithelial, the ovary most connective, and the spleen most immune (**Fig. 1d**).

Across the adult lifespan, cell density declined pervasively, most markedly in the epithelial compartment (**Fig. 1e**), the functional units of most organs. Closer inspection showed how cell-type composition defines organ-specific functional structures (**Fig. 1f**), from ovarian follicles and pulmonary alveoli to thyroid follicles, colonic crypts, seminiferous tubules, and pancreatic acini, whose age-associated changes are therefore expected to compromise organ physiology.

Organ-specific modeling confirmed declining density with age in 8 of 16 organs (FDR < 0.05; BH-corrected linear regression of cells/mm^2^ on age). The ovary was most affected (∼47% reduction), followed by the lung (∼35%), liver (∼27%), spleen (∼26%), adrenal gland (∼25%), prostate (∼24%), vagina (∼23%), and testis (∼21%) (**Fig. 1g**). This loss was visually apparent between young and old donors (**Fig. 1g, Fig. S2a**): the young ovary displayed a dense stroma and abundant follicles (**Fig. S2b**), whereas the aged ovary showed stromal depletion, follicle loss, and expansion of corpus albicans.

### Multi-modal cell phenotyping exposes divergent epithelial aging trajectories

Given the marked decline of epithelial cells with age (Fig. 1e) and their central role in tissue function, we resolved epithelial populations at higher granularity. We developed a multi-modal phenotyping pipeline integrating nuclear (DINOv2) and cellular (CellViT) features, classical morphometric descriptors of nuclear shape, and text-image similarity scores between histological terms and images (PLIP) to classify epithelial cells into nine subtypes (**Fig. 2a**; see Methods). Dimensionality reduction revealed well-separated subpopulations with organ-specific mixing (**Fig. 2b, Fig. S3a-b**).

**Figure 2:**
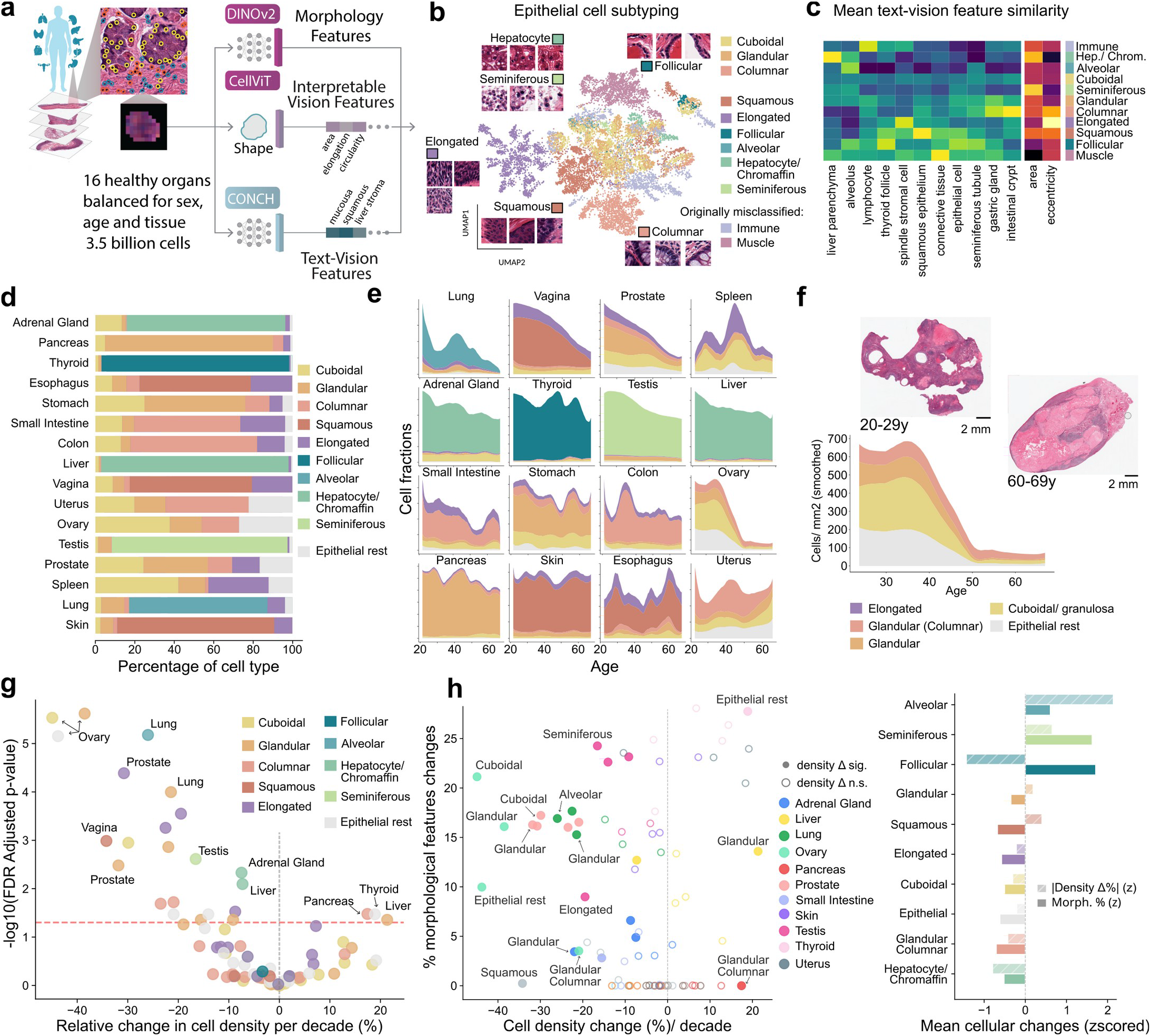
Epithelial subtyping reveals organ-specific composition and age-associated changes in cell density and morphology. **a)** Cell phenotyping pipeline: segmentation and feature engineering with CellViT and DINOv2, integrated with morphological descriptors and vision-text neighborhood queries for epithelial subtype annotation. **b)** UMAP of slide-level mean fused features (CellViT + DINOv2, averaged per slide and epithelial subtype). **c)** Heatmap of mean CONCH/PLIP text-image similarity scores and nuclear morphology features per epithelial cluster, used for annotation (topmost specific terms shown). **d)** Epithelial subtype composition across organs (stacked proportions per organ). **e)** Epithelial subtype densities across age per organ (stacked area; 15-sample rolling mean and Savitzky-Golay smoothing). **f)** Epithelial subtype densities across age in the ovary (as in e), with representative young and old donor WSIs. **g)** Age-associated changes in epithelial subtype densities across organs (OLS on log1p densities; covariates, sex and ischemic time; dashed line, FDR-adjusted p = 0.05). **h)** Age-associated density change versus morphological remodeling. Left: density change per decade versus mean % of significantly changing DINOv2 and CellViT features per cell-type-organ combination (filled, significant density change at FDR < 0.05). Right: composite aging scores ranked by cell type.

Annotating clusters through text-image similarity (**Fig. 2c**) resolved nine subtypes spanning two categories: ontogenically defined, organ-specific populations (hepatocytes/chromaffin, seminiferous, follicular, squamous, and alveolar cells), and morphologically defined classes shared across organs (cuboidal, glandular, glandular columnar, and elongated cells) (**Fig. S3c**). The elongated class was morphologically rather than ontogenically driven, comprising columnar epithelium in gastrointestinal tissues, squamous epithelium in vagina and skin, and stromal cells within the epithelial compartment in the female reproductive system. This strategy also reclassified cells originally misassigned by CellViT as epithelial or neoplastic, including immune and muscle cells. The resulting subtype composition captured known specialization (**Fig. 2d**), with hepatocytes dominating the liver, glandular columnar cells the pancreatic acini, and squamous cells the esophagus.

Quantifying subtype-specific densities across the lifespan revealed pervasive decline in most organs, with occasional fluctuations likely tied to subclinical pathology (**Fig. 2e**). Ovarian aging was most dramatic (**Fig. 2f**), with near-complete depletion of granulosa cells (cuboidal subtype) alongside stromal loss and corpus albicans expansion, refining the organ-level trend (**Fig. 1g**). Systematic testing across all subtype-organ combinations (**Fig. 2g**) confirmed widespread, organ-specific declines. The strongest was ovarian granulosa loss (∼45% per decade, toward near-complete depletion), consistent with follicular loss driven by granulosa apoptosis^27–29^. Alveolar cells in the lung (∼25-30% per decade) tracked documented type 2 alveolar loss and pro-fibrotic remodeling^30,31^, and prostatic glandular cells (∼25-30% per decade) the near-linear decline of glandular volume fraction despite stromal expansion in benign prostatic hyperplasia^32^. Seminiferous cells in the testis fell ∼20% per decade, consistent with germ cell loss and tubule involution^33^. The only increases were pancreatic glandular columnar cells (+17%/decade, FDR = 0.033), consistent with ductal changes including PanIN accumulation and ductal ectasia^34^; and two others which likely reflect morphological drift rather than true expansion (thyroid residual epithelium, +19%/decade; liver glandular cells, +21%/decade), given the purely morphology-based phenotyping.

Besides cell abundance, aging also reshaped cell morphology. In the vagina, morphological embeddings revealed an age-dependent gradient from distinct young clusters toward a mixed older phenotype (**Fig. S3d**), indicating erosion of morphological distinctions between subtypes. Systematic modeling of per-cell-type feature spaces identified the uterus as most affected, with all three broad populations remodeling substantially (**Fig. S3f**), consistent with documented disruption of its epithelial, stromal, and immune compartments^35,36^. Jointly quantifying density and morphology (**Fig. 2h**) showed many populations undergo concurrent loss and transformation: the prostate combined 20-32% per-decade declines across all subtypes with ∼16-17% feature remodeling, and the ovary the largest density loss (granulosa, -45%) with a ∼21% morphological shift, whereas thyroid subtypes showed the inverse, stable densities but the highest remodeling (25-28% of features), consistent with follicular involution, cystic atrophy, and colloid depletion^37^. Composite aging scores (**Fig. 2h**) ranked alveolar cells as most density-affected and follicular cells as most stable, with seminiferous and follicular cells showing the strongest morphological remodeling.

This resolved morphological phenotyping offers a quantitative view of age-associated epithelial and stromal cell dynamics across human organs, and to our knowledge, the first systematic characterization of cell-centric age-associated remodeling at this scale for many organs, such as the uterus.

### Morphological subtypes are molecularly coherent and organ-specific

Because our cell types, including the nine epithelial subtypes, were defined from H&E morphology alone, we asked whether they correspond to transcriptionally coherent populations. Leveraging matched GTEx bulk RNA-seq, we modeled per-slide cell-type densities against gene expression using an organ-stratified negative-binomial GLM with slide area as an offset and age, sex, and ischemic time as covariates, treating each organ and cell-type pair independently across all morphological cell types (**Fig. 3a**).

**Figure 3:**
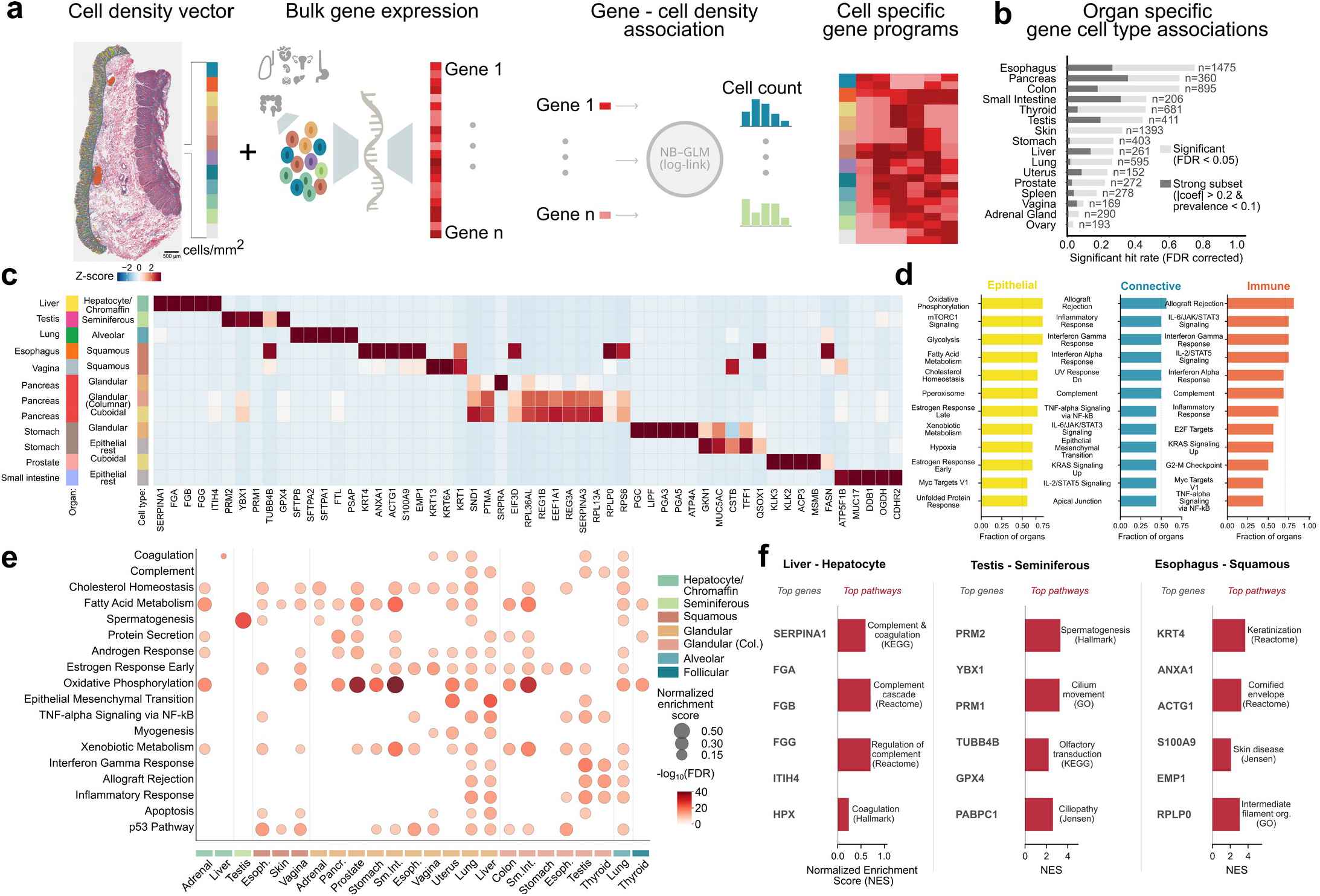
Morphological cell types are molecularly coherent and organ-specific. **a)** Illustration of the analytical strategy where per-slide cell-type densities are modeled against gene expression via an organ-stratified negative-binomial model with donor co-variates, yielding cell-type-specific gene programs. **b)** Fraction of significant genes associated with cell types at Benjamini-Hochberg FDR < 0.05. **c)** Top positively associated genes across cell types and organs. **d)** Most recurrent MSigDB Hallmark pathways associated across broad cell type lineages. **e)** Top positively enriched Hallmark pathways for parenchymal cell types across organs. **f)** Examples of top associated genes (ranked by coef. × SD) and enriched pathways in liver hepatocytes (complement and coagulation), testis seminiferous cells (spermatogenesis and cilia), and esophagus squamous cells (keratinization).

Across 16 organs, 392,103 gene associations were significant (FDR < 0.05; **Fig. 3b, Fig. S4a**). Genes tied to a given morphological label were almost entirely non-overlapping across organs (median Jaccard below 0.02), confirming that these labels capture robust, organ-specific transcriptional identities (**Fig. 3c, Fig. S4b**).

The same morphological class resolved distinct abundance-associated programs in different organs (**Fig. 3c**). Squamous cells were marked by keratin expression (KRT1/4/6A/13), glandular cells showed digestive and acid-secretory genes in the stomach (LIPF, PGA3/5), consistent with gastric chief and parietal cells respectively, but protein-synthesis and protein-targeting machinery in the pancreas (SRPRA, SND1, EEF1A1). Within the stomach, general epithelial cells resolved a distinct gastric mucous program (MUC5AC) consistent with foveolar cells. Organ-restricted subtypes recovered their defining identities: fibrinogen, and protease and endopeptidase inhibitor production in the liver (FGA/B/G, SERPINA1, ITH4), and alveolar type II surfactant proteins in the lung (SFTPA/B). These signatures extended to the pathway level, where connective lineages shared a fibro-inflammatory program, immune lineages a pan-immune program, and epithelial lineages the most organ-diverse functions, resolving steroidogenic versus citrate-accumulating metabolism in adrenal versus prostatic glandular cells despite a shared morphology (**Fig. 3d-e**). Among others, this convergence was clearest for liver hepatocytes, testis seminiferous cells, and esophageal squamous cells, where top genes and multi-library enrichments agreed (**Fig. 3f**).

These analyses provide a molecular grounding for the histologically discovered epithelial subtypes, showing that populations defined from morphology alone are transcriptionally coherent and organ-specific, though inferred from bulk rather than spatially resolved expression.

### Gastrointestinal vascular architecture is organized along transmural gradients that decline with age

Having established that individual cell types are molecularly coherent, we turned from cellular identity to the spatial architecture into which cells assemble, beginning with the vasculature that sustains nutrient absorption, immune trafficking, and tissue homeostasis in the gastrointestinal tract, and whose decline is a key substrate of age-related dysfunction. Detecting endothelial cells with a classifier trained on ∼27,000 annotations and assembling them into ∼65,000 vessels represented by CONCH morphology, we resolved six phenotypes spanning the vascular hierarchy: arteries, arterioles, capillaries, intramuscular vessels, veins/lymphatics (grouped as morphologically indistinguishable), and blood-filled veins (**Fig. 4a-d, Fig. S5a**). As for epithelial subtypes, vessel identity rather than organ of origin drove the dominant axis of morphological variation (**Fig. 4e**), indicating a vascular vocabulary shared across the gastrointestinal tract.

**Figure 4:**
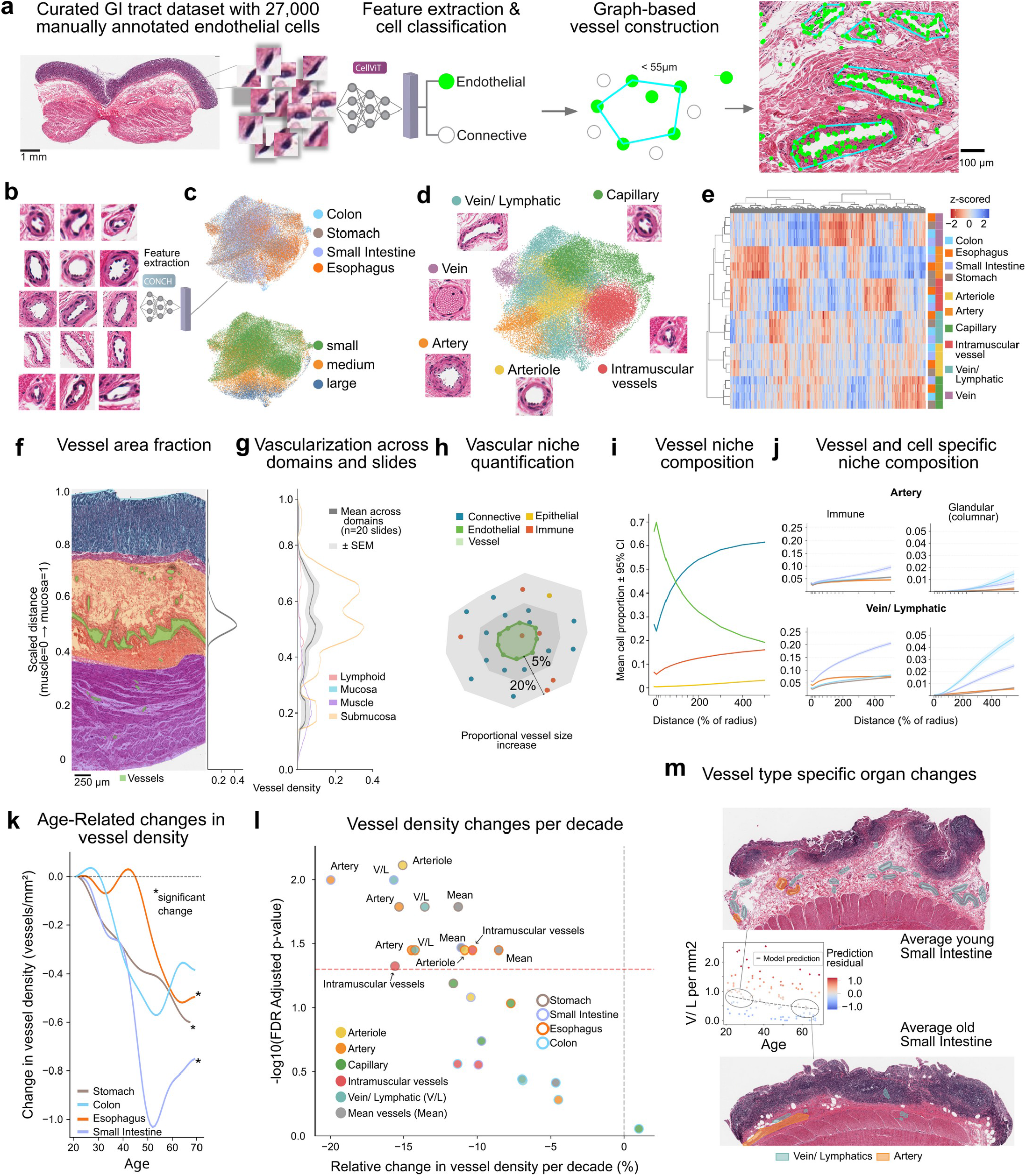
Vascular architecture reveals organ-specific vessel composition and age-associated density loss. **a)** Vessel analysis pipeline, from endothelial cell annotation and classification to graph-based vessel construction, with example segmentation overlaid on a tissue crop. **b)** Example vessel crops used for CONCH feature extraction. **c)** UMAP of vessel-level CONCH embeddings colored by organ (top) and vessel area group (small, medium, large; bottom). **d)** The same UMAP colored by annotated vessel subtype. **e)** Clustermap of mean CONCH features per vessel type and organ. **f)** Example section with overlaid domain and vessel annotations (colored as in j), with its transmural vessel area fraction gradient. **g)** Transmural vessel area fraction gradient for stomach (n = 10) and small intestine (n = 10), with domain-specific contributions. **h)** Vascular niche analysis: concentric radial buffers around a vessel quantify cell type proportions with distance. **i)** Cell type composition gradients around vessels across all organs and vessel types, for epithelial cell types (mean, 95% CI). **j)** Vessel-type-specific niche gradients for arteries (top) and veins/lymphatics (bottom), for immune and glandular columnar cells. **k)** Smoothed total vessel density across age per organ; significance from mean vessel density Gamma GLMs (as in l). **l)** Age-associated vessel subtype density changes per decade across organs (Gamma GLM, organ-level FDR-corrected). **m)** Vein/lymphatic density versus age in the small intestine (Gamma GLM trend), with representative young and old small intestine crops and manual vessel annotations.

Vessel density reflected organ physiology, being highest in the absorptive small intestine and esophagus (median 2.03 and 1.66 vessels/mm^2^) and lower in the colon and stomach (1.31 and 0.95 vessels/mm^2^) (**Fig. S5b-c**). Veins and lymphatics dominated every organ and were especially rich in the small intestine, while intramuscular vessels and capillaries were most abundant in the muscularis-rich esophagus, and arterioles consistently outnumbered arteries (**Fig. S5b**).

The wall itself imposed a stereotyped spatial order on this vasculature. Quantifying vessel area fraction along the transmural axis from mucosa to muscle revealed a near-avascular mucosa, a first peak at the muscularis mucosae, and a dominant peak in the submucosa, consistent with its role in nutrient supply and lymphatic drainage to the mucosa and its tertiary lymphoid structures^38–40^ (**Fig. 4f-g**). Submucosal vascularization was itself bimodal, peaking near both the mucosa and the muscle, and a further peak arose at the center of the muscularis where vessels perfuse the smooth muscle. This architecture extended to the cells surrounding each vessel: the immediate perivascular niche was dominated by endothelial cells (∼70%) with connective cells (∼25%) and few immune cells (∼5%), giving way with distance to connective cells of the perivascular smooth muscle and stroma, and to a steadily rising immune fraction (∼15% at the periphery), with arteries and veins/lymphatics occupying distinct niches (**Fig. 4h-j**).

Against this conserved baseline, aging thinned the vasculature. Total vessel density declined with age in every organ, significantly so in the small intestine, stomach, and esophagus (**Fig. 4k**), and the small intestine was most vulnerable, losing its arteries fastest (20% per decade) followed by veins/lymphatics (15%) (**Fig. 4l**). This rarefaction was visually stark in the small intestine (**Fig. 4m**), where a richly vascularized young submucosa gave way to a stiffened, fibrotic, vessel-poor submucosa in older donors, consistent with the microvascular rarefaction of aged tissues^41–43^. Capillary-scale vessels are likely underrepresented, as structures with fewer than six endothelial cells were filtered for robustness, and erythrocyte-filled veins were excluded from age analyses for insufficient per-bracket sampling.

In these analysis, the resolved gastrointestinal vasculature emerged as a conserved system organized along stereotyped transmural gradients and vessel-type-specific niches. The large-scale quantification highlighted a pervasive age-associated rarefaction most pronounced in the arteries of the small intestine.

### Functional tissue units remodel with age beyond what cell composition predicts

Tissue function depends not only on cell abundance but on how cells assemble into functional units such as crypts, follicles, and lobules. To capture this, we built cell spatial proximity graphs per slide (**Fig. 5a**, step 1), whose topology separated organs by architectural identity (step 2), with cell-dense organs, gastrointestinal tissues, and skin forming distinct groups, and within each organ samples drifted with age toward a more similar topology (step3), consistent with the previously observed pervasive density loss (**Fig. 1e-g**). Community detection then partitioned each section into spatially coherent units, which we clustered per organ by composition and topology into archetypes we term functional tissue units (FTUs)^1,44^ (step 4-5). Projected back onto tissue, these corresponded to recognizable compartments such as crypts, stroma, and vascular niches (step 6).

**Figure 5:**
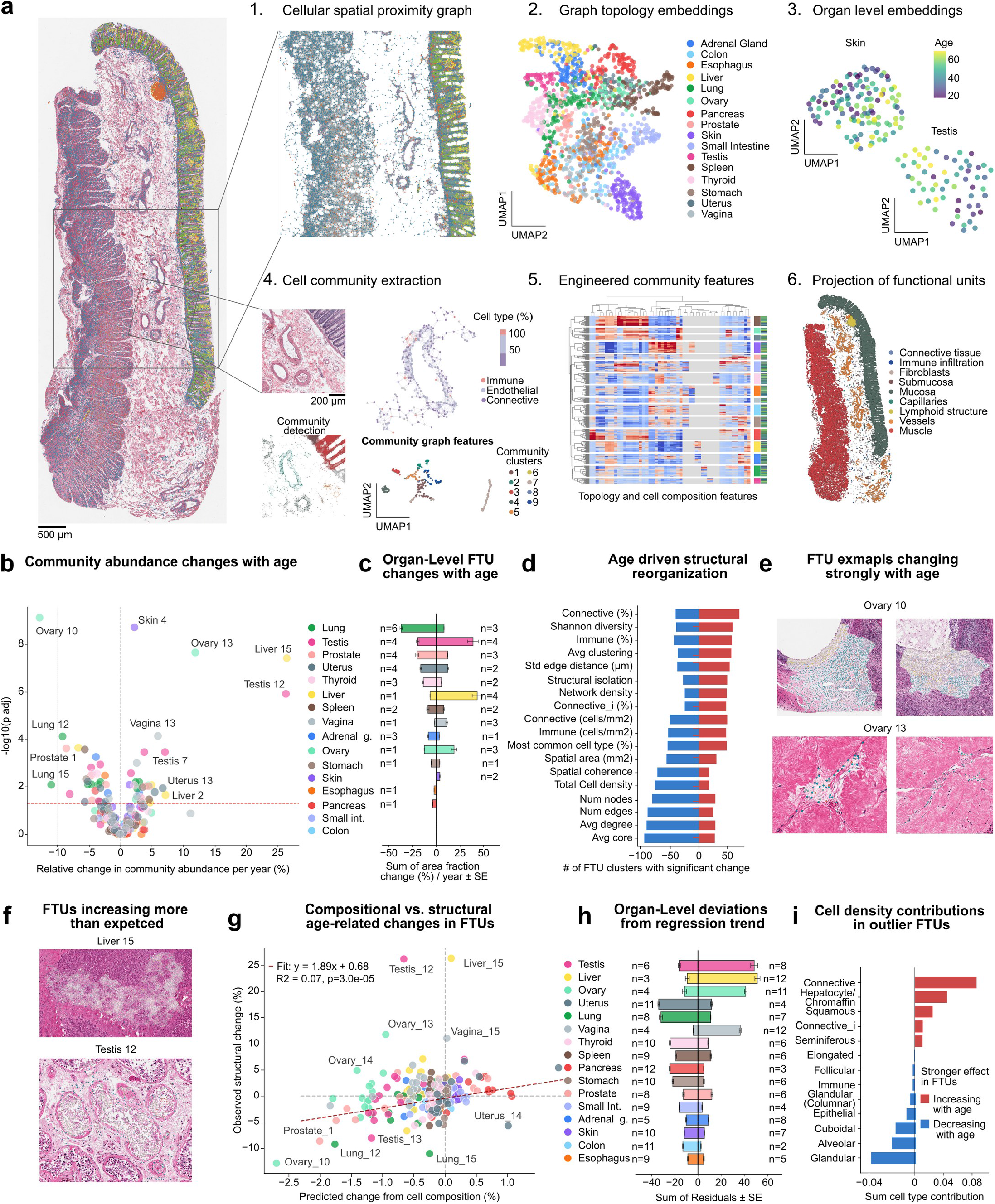
Functional tissue units change in abundance and composition with age, beyond cell density shifts. **a)** Illustration of the process of detection and characterization of FTUs in whole slide images: spatial proximity graphs are built with edges connect cells within 50 μm (1); slide-level cell graph topology features are extracted and can be characterized across organs (2) and within (3); cell communities are extracted via Leiden clustering (4) and can be characterized by cell type composition or community-level features; finally, recurrent cell communities within each slide are jointly clustered into FTU meta communities (5) and projected back in space for characterization (6). **b)** FTU abundance change with age (Gamma GLM): percentage change per year versus significance (local FDR), colored by organ (as in c, throughout). **c)** Per-organ effect sizes of significant FTU changes, sorted by number of changing units. **d)** The 18 most frequently changing community features, by frequency of significant increase or decrease across FTUs (local FDR < 0.05). **e)** Exemplary age-affected ovarian FTUs. **f)** Exemplary FTUs expanding faster than expected from composition. **g)** Observed FTU abundance change versus that expected from compositional change, per year, with linear fit. **h)** Per-organ directional regression residuals, sorted by summed absolute residual, marking organs where FTU change exceeds compositional expectation. **i)** Difference in mean absolute compositional contribution per cell type between FTUs expanding versus declining more than expected from composition (top cell types by magnitude).

Testing each FTU for age-associated abundance change (Gamma GLM; see Methods), 62 of 236 FTUs across 16 organs changed significantly (26%, FDR < 0.05), balanced between gains (30) and losses (32) (**Fig. 5b-c, Fig. S5b**). Remodeling was highly organ-specific, most extensive in the lung, testis, prostate, and uterus and undetectable in the colon and small intestine (**Fig. 5d**). Across affected FTUs, a coherent signature emerged (**Fig. 5d**): features of structural densification declined most (cell density -62%, spatial coherence -60%, edges -57%, nodes -48%, average degree -52%), while diversification and loosening increased (connective +27%, immune +14%, Shannon diversity +19%, structural isolation +28%). Individual units traced this same trajectory, from the steep loss of granulosa-rich ovarian follicles (Ovary_10: -12.9%/year, p_adj = 7.4 × 10^−10^) and their replacement by stromal units in corpus albicans (Ovary_13: +11.8%/year, p_adj = 2.2 × 10^−8^) (**Fig. 5e**) to fibrotic liver expansion (Liver_15: +26%/year) and sclerotic seminiferous atrophy (**Fig. 5f**), alongside known changes such as epidermal thinning^45^ and alveolar loss^46^ (**Fig. S6a**). Tissue aging thus follows a shared trajectory from dense, specialized units toward sparser, more heterogeneous structures enriched in stromal and immune cells, recurring across organs despite their distinct cellular constituents.

Critically, this trajectory was not a byproduct of the cell-composition changes documented above. Comparing each FTU’s observed abundance change to that expected from the age trajectories of its constituent cell types (**Fig. 5g**), composition explained only 7% of variance (R^2^ = 0.07, p = 3.0 × 10^−5^), establishing that FTU remodeling is largely decoupled from cell composition. This decoupling was greatest in the testis, liver, and ovary (**Fig. 5h**), and at the cell-type level, glandular, alveolar, and cuboidal cells marked units declining beyond expectation while connective cells, hepatocytes, and squamous cells marked those expanding beyond it (**Fig. 5i**). This coordinated loss of specialized parenchymal units and gain of stromal or fibrotic compartments is an organizational mode of aging that exceeds the sum of individual cell density shifts and is most pronounced in the most specialized organs.

### Spatially compartmentalized immune and stromal remodeling resolves gastrointestinal inflammatory pathology

The framework developed above also generalizes to pathological remodeling. Leveraging GTEx subclinical pathology annotations, we compared healthy stomach (n = 112) and esophagus (n = 35) slides with slides annotated for gastritis, esophagitis, or inflammation (stomach n = 54, esophagus n = 41; **Fig. 6a, Fig. S7a**), a set held out from the aging analyses above.

**Figure 6:**
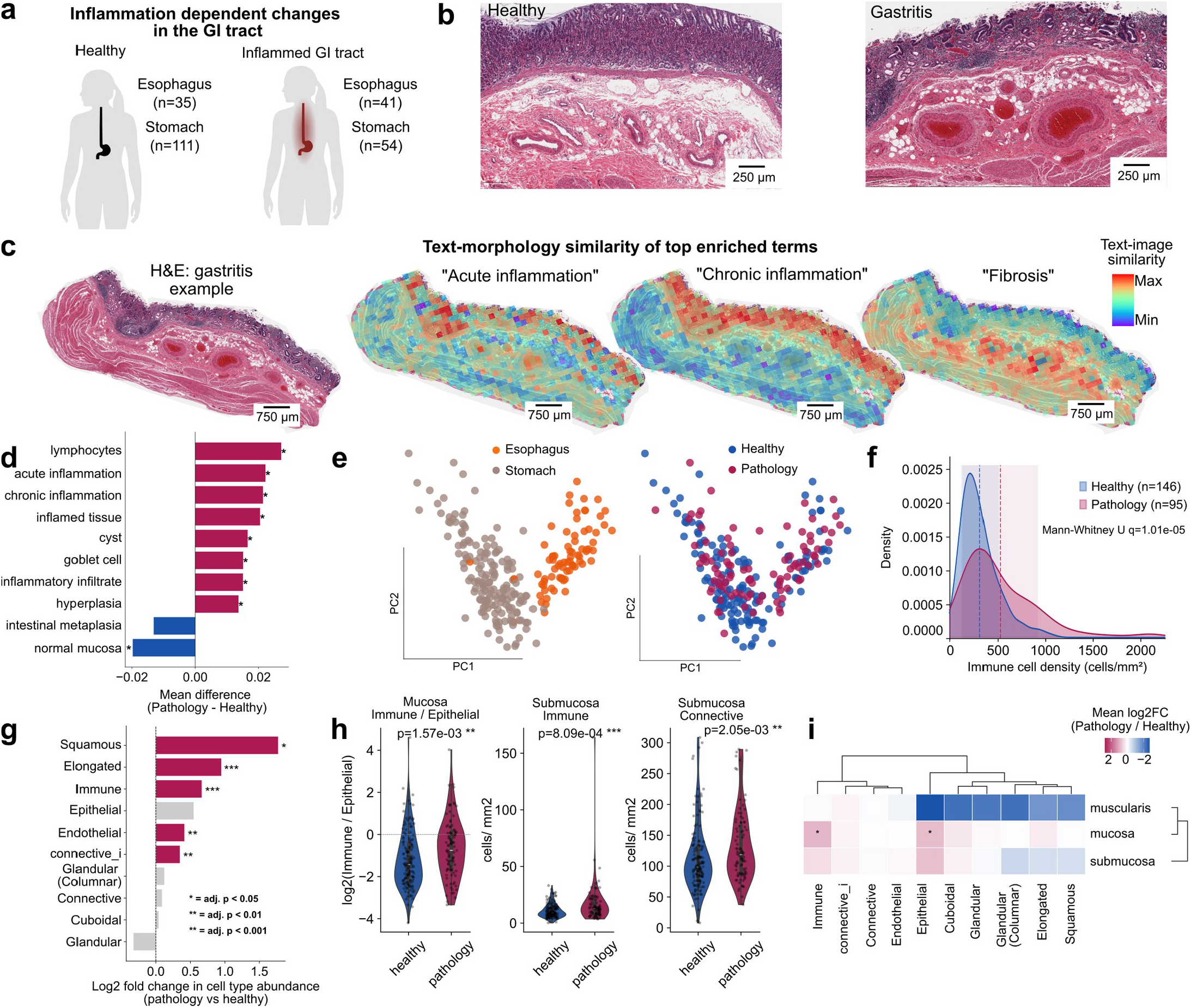
Gastrointestinal pathology shows domain-specific immune infiltration and morphological remodeling. **a)** Sample distribution. **b)** Example healthy versus gastritis stomach WSIs displaying mucosa and submucosa. **c)** Representative gastritis WSI with tile-level CONCH zero-shot similarity scores for acute inflammation, chronic inflammation, and fibrosis. **d)** Top 10 CONCH text-image similarity terms by mean difference between pathology and healthy slides (Wilcoxon rank-sum, BH-corrected; * adjusted p < 0.05). **e)** UMAP of WSIs by cell type densities, colored by organ and pathology status. **f)** Immune cell density in healthy versus pathology slides, both organs combined (kernel density estimate; dashed lines, means; bands, ±1 SD; Mann-Whitney U, BH-corrected). **g)** Log2 fold change of cell type densities between pathology and healthy slides across organs. **h)** Immune-to-epithelial ratio in the mucosa (left) and submucosal cell type densities (right), healthy versus pathology, across shared domains (Mann-Whitney U). **i)** Mean log2 fold change of domain-specific cell type densities between pathology and healthy slides, averaged across stomach and esophagus (shared domains, mucosa aggregated; Mann-Whitney U, BH-corrected).

Representative stomach sections illustrate the tissue-level hallmarks of gastritis (**Fig. 6b**): healthy mucosa shows well-organized gastric glands and foveolar epithelium over a loose, well-vascularized submucosa, whereas gastritis disrupts this architecture with dense immune infiltration, tertiary lymphoid structures, and a fibrotic submucosa with perivascular fibrosis. To resolve these alterations without supervision, we applied CONCH zero-shot text-image similarity scoring (**Fig. 6c, Fig. S7b**). In a representative gastritis slide, acute inflammation scored highest at mucosal immune infiltrates and tertiary lymphoid structures, chronic inflammation was broader across the mucosa, and fibrosis localized to the submucosa. Across the cohort, mean similarity scores discriminated pathology from healthy tissue at the slide level, enriching lymphocytic, inflammatory, and hyperplasia terms in pathology and normal mucosa in healthy slides (**Fig. 6d**).

Cellular composition was also distinct (**Fig. 6e**). Immune cell density was significantly elevated in both organs, with healthy slides peaking sharply near 250 cells/mm^2^ (SD 193) and pathology slides peaking near 300 cells/mm^2^ with a far wider distribution (SD 391; Mann-Whitney U p = 1.01 × 10 ^-5^), reflecting heterogeneous infiltration (**Fig. 6f**). Remodeling extended beyond the immune compartment (**Fig. 6g**): squamous epithelial cells showed the largest change (log_2_ FC = 1.77), consistent with basal cell hyperplasia in chronic inflammation ^47^, followed by elongated stromal cells (0.95), immune cells (0.66), and endothelial cells (0.41). The residual epithelial class also rose, consistent with inflammation-induced erosion of normal epithelial identity, a feature rather than a limitation of the approach. Mapping these changes onto microanatomical domains revealed a consistent spatial pattern across both organs (**Fig. 6h-i**): the mucosa showed an elevated immune-to-epithelial ratio (p = 1.57 × 10^−3^), the submucosa coordinated immune and connective expansion consistent with inflammatory infiltration and stromal remodeling, and the muscularis was largely spared.

Together, these results illustrate how the same framework that resolves physiological aging also resolves inflammatory disease, integrating single-cell phenotyping, microanatomical domain mapping, and vision-language scoring into a spatially explicit, interpretable view of tissue remodeling.

## Discussion

The cellular and molecular hallmarks of aging are well established, but how they accumulate into the architectural changes that compromise organ function has been difficult to resolve at scale. By analyzing the histomorphology of 3.5 billion cells across 16 human organs and the adult lifespan, we show that cell density, morphology, vascular architecture, and tissue organization remodel with age along shared principles but organ-specific manifestations, positioning histopathology as a population-scale, lifespan-resolved resource for spatial biology.

The most distinctive finding is that the architectural layer of aging is largely decoupled from cell composition (R^2^ = 0.07). Over a quarter of functional tissue units changed with age along a coherent trajectory, from dense, specialized units toward sparser, more heterogeneous structures enriched in stromal and immune cells, recurring across organs as diverse as the lung, testis, prostate, and uterus despite their distinct cellular constituents. This is consistent with the progressive replacement of functional parenchyma by stromal and immune compartments that characterizes age-associated fibrosis, chronic low-grade inflammation, and declining regenerative capacity^10^. That the organs with the greatest unexplained remodeling are also the most parenchymally specialized suggests aging erodes precisely the organizational features that make a tissue specialized. Cell counts, even stratified by type, therefore capture only one axis of aging, one invisible to dissociative single-cell methods but resolved by population-scale histopathology.

Beyond morphology, pairing histology with matched bulk transcriptomes showed that cell types defined from H&E alone are transcriptionally coherent and organ-specific. This validates the atlas, but also illustrates a broader principle. Just as histopathology at scale resolves spatial detail across whole cross-sections and the full lifespan that spatial transcriptomics cannot yet reach, associating cell densities with bulk RNA-seq across many individuals recovers cell-type-resolved molecular signal of the kind usually accessible only to single-cell RNA-seq, without dissociation. This adds a molecular dimension to the vast archives of legacy histology accumulated in clinical and biobanking settings, and we expect it to deepen considerably with spatially resolved data in future work.

This work resolves the cellular layer of human tissue aging, complementing our previous characterizations of slide-level aging signatures^24^ and microanatomical remodeling^25^. Where that work operated at the tile level (smallest unit ∼120 µm), here we resolve individual cells, vessels, and tissue units, together establishing a multi-scale framework for population-scale spatial biology that complements molecular cellular atlases^14,15^ at a scale and life-span coverage molecular methods cannot yet reach. Its extension to gastrointestinal inflammatory pathology, where interpretable vision-language scoring spatially resolved acute inflammation, chronic inflammation, and fibrosis, further shows the framework generalizes from physiological aging to disease, and is well-suited to retrospective interrogation of existing histopathology archives.

Several limitations apply. The GTEx cohort is postmortem, rapid-autopsy collected, cross-sectional, with variable ischemic times and a 1:2 female-to-male ratio; ischemic time is modeled throughout, but autolytic effects cannot be excluded, causal ordering is inaccessible, and sex-specific dynamics are underpowered. Cell, vessel, and FTU representations depend on fixed spatial scales, and the six-cell minimum for vessel construction likely underrepresents capillaries. CellViT was trained predominantly on cancer tissue, a bias mitigated after inference through multi-modal phenotyping. Finally, cell and vessel subtypes rest on morphology and vision-language features rather than molecular markers, so some, such as the elongated class or the combined vein/lymphatic label, are structurally rather than ontogenically defined.

This work establishes the cellular and architectural trajectory of tissue aging as a quantitative phenotype measurable retrospectively, at population scale, from the most widely available tissue assay. The atlas of 3.5 billion cells, ∼65,000 vessels, and 236 functional tissue unit types is a foundational reference against which future longitudinal, interventional, molecular, or spatial studies can be anchored to resolve the mechanisms of architectural decline in the aging human body.

## Methods

### Data reporting

No statistical methods were used to predetermine sample size due to the predetermined sample availability of the GTEx project^48^. From the GTEx cohort, 16 organs (20 tissue subtypes) were selected. Cell segmentation, feature extraction and classification were performed on all available WSIs across the 16 organs (14,788 WSIs from 980 donors). A balanced dataset was constructed by sampling 12 slides per age bracket (20-29, 30-39, 40-49, 50-59, 60-69), sex and organ combination, yielding 120 slides per organ for organs with both sexes represented. For sex-specific organs (ovary, prostate, testis, uterus, vagina), 12 slides per age bracket were sampled from the available sex, yielding 60 slides per organ. Where fewer than 12 slides were available for a given stratum, the maximum available number was used (adrenal gland: 117, liver: 115, uterus: 59, vagina: 59). The balanced dataset comprised 1,610 WSIs from 646 donors. 25 slides were excluded from cross-organ analyses after manual inspection revealed incorrect tissue annotations. The balanced dataset was used for epithelial subtype discovery and all downstream statistical age-associated analyses. One colon slide (GTEX-WHSE-2626) was excluded due to severe autolysis. An additional 88 pathology-annotated slides (46 stomach, 42 esophagus) from 83 donors were included for the gastrointestinal pathology comparison but held out from all age-associated analyses.

### Data acquisition

Whole-slide images from the GTEx project were obtained through the publicly available GTEx data portal. Image digitization was previously performed using a Leica Biosystems Aperio ScanScope at 20× magnification, yielding an effective resolution of 0.4942 μm per pixel. Associated metadata, comprising precise chronological age, demographic characteristics, lifestyle factors, and clinical attributes of donors, along with transcriptomic data were retrieved from dbGaP, as they are subject to controlled-access restrictions.

### Cell segmentation and classification

CellViT with the SAM-H backbone (CellViT-SAM-H-x20) was used to perform cell segmentation and classification on H&E-stained WSIs. Slides were preprocessed by extracting patches of 1,024 × 1,024 pixels at 20x magnification with 6.25% overlap, a minimum intersection ratio of 0.05 and Macenko stain normalization, using the cucim backend. Cell detection was performed on the extracted patches using a single NVIDIA RTX A6000 GPU. CellViT outputs per-cell annotations including nuclear contours, centroids and classification into five broad cell types (epithelial/neoplastic, inflammatory, connective, dead and neoplastic). Cells classified as ‘dead’ were excluded from all downstream analyses. The remaining four cell types were carried forward, with neoplastic cells merged into the epithelial class.

### Nuclear feature extraction

For each segmented nucleus, a 56 × 56-pixel region centered on the nuclear centroid was extracted from the WSI at full resolution (level 0, 20x, ∼0.499μm). Within this region, a binary mask was constructed from the CellViT-derived nuclear contour polygon using OpenCV^49^ and applied to zero out all extranuclear pixels, isolating the nuclear content. Masked nuclear images were resized to 224 × 224 pixels using bicubic interpolation and normalized using ImageNet statistics (mean = [0.485, 0.456, 0.406], s.d. = [0.229, 0.224, 0.225]). Feature extraction was performed using a frozen DINOv2 ViT-S/14 model (dinov2_vits14, loaded via torch.hub). The resulting 384-dimensional CLS token embedding was used as the feature representation for each nucleus. All nuclei were processed in batches of 1,024 on HPC CPUs.

Classical morphological features were computed directly from the CellViT-derived nuclear contour polygons. For each slide, a shared integer-labeled mask was constructed by filling all nuclear contour polygons onto a common coordinate space using OpenCV^49^, where each nucleus received a unique integer label. Morphological descriptors were then computed jointly using scikit-image’s regionprops^50^55, including area, bounding box area, convex area, filled area, major and minor axis lengths, eccentricity, equivalent diameter, Euler number, extent, maximum Feret diameter, perimeter, Crofton perimeter, solidity, inertia tensor and eigenvalues, and image moments (central, normalized and Hu). From these, additional derived features were calculated: elongation (ratio of major to minor axis length) and circularity (4*pi * area / perimeter^2^). Contour curvature statistics were also computed following the approach described in sc-MTOP^51^, including mean, standard deviation, maximum and minimum curvature, as well as normalized counts of protrusions and indentations identified as local extrema along the contour curvature profile within a sliding window of size 5.

### PLIP cell neighborhood feature extraction

To characterize the histological microenvironment surrounding each segmented cell, we computed text-image similarity scores between cell-centered tile neighborhoods and a curated panel of histopathology text terms using the PLIP^22^ vision-language model. LazySlide^52^ was used to detect tissue regions, tile tissue into 224-pixel patches at ∼0.5 μm/pixel, extract PLIP tile features, and compute cosine similarities between PLIP text embeddings of a term panel describing epithelial subtypes and stromal/structural reference categories and per-tile image embeddings, yielding a per-tile term-similarity matrix per slide.

Per-cell similarities were obtained by spatially joining cell centroids (from the CellViT-derived segmentation) to tile polygons using geopandas^53^, assigning each cell to the containing tile via a within predicate, with boundary duplicates kept once. Each cell thus inherited the similarity vector of its parent tile. Per-slide matrices were concatenated and restricted to the cells retained in the fused, filtered cell AnnData^54^.

### Epithelial subtyping

To identify epithelial subtypes, up to 2,000 epithelial cells (or the maximum available, including neoplastic-classified cells) were randomly sampled per WSI from the balanced dataset, excluding cells within 128 pixels of the patch border, as CellViT’s Vision Transformer-based feature extraction yields less reliable cell representations near patch boundaries due to incomplete spatial context and potential cell truncation. CellViT and DINOv2 features were combined into a unified cell-level representation via PCA-based feature whitening ^55^. PCA with whitening was applied independently to the CellViT and DINOv2 embeddings, each retaining 50 components. The resulting whitened features were concatenated into a single 100-dimensional representation per cell. Leiden clustering (igraph flavor, resolution 0.7) was performed on the fused feature space. Cell image crops of the most central cells per cluster (closest to the cluster centroid in embedding space) were extracted from the WSIs using the CellViT contour polygons for visual validation. Three clusters were identified as segmentation artifacts and removed. The remaining cells were re-embedded (PCA, neighbor graph, UMAP) and reclustered using Leiden (igraph flavor) at multiple resolutions including 1.5, which was used for downstream annotation.

Cluster annotations were determined by jointly inspecting the fused-feature cluster structure, the classical nuclear morphology features (computed via scikit-image regionprops, as described above) and PLIP^22^ vision-text similarity scores. PLIP text similarities were computed for each cell by assigning it to the nearest WSI tile (by Euclidean distance to tile centroids) and transferring the tile-level similarity values from a curated set of cell type terms. Final cell type labels were assigned through manual curation of the cluster annotations at resolution 1.5.

To predict epithelial subtypes for all epithelial cells across all 14,788 WSIs, a multi-layer perceptron (MLP) classifier was trained on the CellViT features using the manually curated cluster labels. Additional artifact clusters identified during annotation were excluded from the training set. The MLP consisted of two hidden layers (512 and 128 units) with batch normalization, ReLU activation and dropout (p = 0.2) after each hidden layer, followed by a linear output layer. The model was trained using the Adam optimizer (learning rate = 1 × 10^−3^) with balanced cross-entropy loss (class weights = N / (K * n_c)) and mixed-precision training. A 90/10 train/validation split was performed stratified by slide and cell type label. Early stopping with a patience of 5 epochs was applied based on the validation loss, and the model checkpoint with the lowest validation loss was retained.

### Age-associated changes in cell type density and morphology

To assess age-related changes in cell composition and nuclear morphology across tissues, we modeled the relationship between donor age and two sets of cell-level summary statistics per tissue sample. For cell type density, the number of cells of each type per slide was normalized by the tissue area (computed from the microns-per-pixel resolution of each WSI) to obtain cell densities. Density values were log-transformed (log1p) and modeled using OLS regression with age as the primary predictor and sex and ischemic time as covariates. The percentage change per year was derived from the regression coefficient as (exp(β) − 1) × 100. P-values were corrected using the Benjamini-Hochberg FDR procedure across organs within each cell type and corrected globally across all cell type-organ combinations.

To identify age-related changes in nuclear morphology, we analyzed the relationship between donor age and cell-level feature representations aggregated per cell type and WSI. For each slide, CellViT and DINOv2 embeddings were separately averaged across all cells of a given type, yielding one mean feature vector per cell type per slide. Each feature was then modeled independently on its original scale using OLS regression with the same covariates. The percentage change per year was calculated as 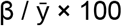, where 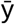 is the mean feature value. P-values were corrected using Benjamini-Hochberg FDR within each cell type-organ combination across features and additionally corrected globally across all cell type-organ-feature combinations. Results were considered significant at an adjusted p-value threshold of 0.05.

### Gene expression association

To associate cell type densities with gene expression, we fit negative binomial generalized linear models (GLM) with a log link function, stratified by organ. For each (organ, cell type, gene) combination, gene counts were modeled as a function of cell type density, with log(slide area) as an offset, and donor age, sex, and ischemic time as covariates. The dispersion parameter was estimated from the null model (count ∼ 1 + covariates + offset) via maximum likelihood and held fixed across all per-gene fits within each organ. Models were fit using statsmodels^56^, and p-values were adjusted within each (organ, cell type) stratum via the Benjamini-Hochberg procedure (FDR < 0.05). Associations passing additional effect-size filters (|coefficient| > 0.2, gene prevalence < 0.1) were designated as strong hits, where gene prevalence was defined as the fraction of (organ, cell type) pairs in which a gene’s expression was non-zero. Per-organ hit rates were computed as the fraction of tested gene x cell-type pairs significant at FDR < 0.05, with strong hit rates computed analogously from the filtered subset.

For parenchymal cell types, we selected the top-5 positively-associated strong genes per (cell type, organ) pair by GLM coefficient. To build a non-redundant gene set across cell-type-organ pairs, we employed a greedy selection: rows were processed in a fixed order (single-organ cell types first, then multi-organ contrasts grouped by cell type), and each row claimed its top-5 genes not already claimed by an earlier row. Each gene therefore appears only once, assigned to the pair for which it shows the strongest association. The resulting coefficient matrix (14 rows x 70 genes) was Z-scored per column for cross-gene comparability. A union-mode version preserving the raw perrow top-5 without deduplication was also generated.

Gene set enrichment was performed via preranked GSEA using gseapy against five Enrichr gene set libraries: MSigDB Hallmark 2020^57^, GO Biological Process 2023^58,59^, KEGG 2021 Human^60,61^, Reactome 2022^62^, and Jensen DISEASES Curated 2025^63,64^. For each (organ, cell type) pair with at least one significant gene, all tested genes were ranked by the signed GLM coefficient without a top-N cutoff. Significant enrichment was defined at FDR < 0.05. For the broad-lineage pathway consensus, fine-grained cell types were aggregated into three categories: connective (Connective, Muscle, connective_i; plus Elongated in Ovary and Uterus), immune (Immune), and epithelial (all remaining). Elongated cells were context-dependent: connective in Ovary and Uterus, epithelial elsewhere. For each Hallmark pathway and category, we computed the fraction of organs in which at least 50% of constituent cell types within that category showed significant up-regulation enrichment. Convergence across the other four libraries was assessed by counting how many independently recovered the same pathway-category pair.

### Endothelial cell detection

To identify endothelial cells within the connective tissue cell population, we trained a binary classifier using manually annotated tissue domains from 14 WSIs of the gastrointestinal tract representing multiple tissues. Tissue regions were annotated into domain categories including endothelium, mucosa, muscle, submucosa, submucosal gland, mucosa border, cells in vessel, granulocyte and serosa. Cells were assigned to domains based on spatial overlap between their centroids and the annotated domain polygons. A balanced training set was constructed with endothelial domain cells as the positive class. Negative examples were sampled from mucosa, muscle and submucosa domains in equal proportions, balanced to match the number of endothelial cells and stratified across tissue samples. Additional negative examples were included from cells in vessel, granulocyte, mucosa border and submucosal gland domains, with the latter two capped at 2,000 cells each. A LightGBM classifier^65^ with default hyperparameters was trained on CellViT morphological features to distinguish endothelial from non-endothelial connective tissue cells. The trained classifier was then applied to predict endothelial identity across all connective tissue cells in gastrointestinal slides of the cohort.

### Vessel segmentation algorithm

To segment vessels in whole slide images, we built spatial proximity graphs from endothelial cell centroids. Endothelial cells were retained for analysis if they had a predicted probability ≥ 0.65 given by the LightGBM classifier. A KDTree was used to connect endothelial cells within 55 μm of each other, and connected components of the resulting graph were identified as candidate vessel structures. Leaf nodes (degree ≤ 1) were iteratively removed to prune spurious connections. For each connected component, a shape representation was computed using an adaptive strategy: single cells were represented as circles with a 15 μm radius, pairs as minimum bounding circles, and larger groups as either convex hulls (for compact, circular arrangements with circularity > or elongation < 2.0) or alpha shapes (for elongated structures). Morphological descriptors were computed for each vessel structure, including area, perimeter, circularity, elongation and extent. These parameters were selected after a systematic search over probability thresholds (0.6-0.95), neighbor distances (20-150 μm), graph construction methods (KDTree, DBSCAN, HDBSCAN), and pruning strategies (none, bridge removal, leaf node removal, combined, high-betweenness edge removal), evaluated across five representative slides with annotations. Vessel structures were identified from endothelial cell niches detected within each slide (minimum 6 cells per structure; polygon area < 772,051 μm^2^, to exclude extreme outliers, which were mistakenly merged vessels). To characterize the cellular microenvironment surrounding each vessel, cell type composition was quantified within concentric buffers at 14 radial distances (0-5× the vessel radius) from the vessel boundary.

### Vessel feature extraction and classification

To characterize vessel morphology and classify vessel types, cropped images of segmented vessel structures were processed using CONCH, a vision-language foundation model pretrained on histopathology images and pathology reports. Vessel images were resized to 448 × 448 pixels using bicubic interpolation and normalized using CLIP normalization statistics (mean = [0.481, 0.458, 0.408], s.d. = [0.269, 0.261, 0.276]). For feature extraction, image embeddings were obtained from the CONCH ViT-B/16 encoder without projection or normalization. For vessel type classification, CONCH’s vision-language alignment was used to compute cosine similarity between projected, normalized image embeddings and text embeddings of predefined histological terms. Text labels were tokenized using the CONCH tokenizer and encoded with the text encoder, and similarity scores between each vessel image and all text labels were computed via dot product in the shared embedding space. Images were processed in batches of 64 on GPUs. Each vessel was classified into one of six subtypes-arteriole, artery, capillary, intramuscular vessel, vein/lymphatic, and vein-by applying Leiden clustering (resolution 0.8) to CONCH (ViT-B-16) visual feature embeddings, followed by manual annotation of resulting clusters. Vessels assigned to size-stratified groups (between 6-20 and >20 endothelial cells) were clustered and annotated separately, then merged; artifacts were excluded. Additionally, vessels were assigned to area-based size groups (small, medium, large) using Gaussian mixture model-derived boundaries on polygon area (small: <4,194 µm^2^; medium: 4,194-21,183 µm^2^; large: >21,183 µm^2^). Capillaries and arterioles in the large area group were removed as likely segmentation artifacts from merged adjacent small vessels (<2% and <0.01% of all capillaries and arterioles, respectively). Vessel subtype densities were computed per slide as the count of each subtype divided by the total tissue area (mm^^2^).

### Vascularization gradient across tissues

To quantify the spatial gradient of vascularization across the gastrointestinal wall, tissue domains (mucosa, submucosa, muscle and lymphoid tissue) were manually annotated in 10 stomach and 10 small intestine WSIs using Cytomine^66^). For each slide, an orientation line was manually drawn perpendicular to the mucosal surface to define the transmural axis. All tissue domain annotations and vessel structures were rotated to align the orientation line with the horizontal axis, and the geometry was flipped if necessary to ensure a consistent mucosa-to-muscle ordering along the y-axis. The mucosa baseline was shifted to y = 0. Vessel structures and tissue annotations were filtered to the x-extent of the orientation line to restrict analysis to a consistent region of interest. Vessel area fraction was computed along the transmural axis in 20 μm bins. For each bin, the geometric intersection between the bin rectangle and each tissue domain was calculated, and vessel polygons overlapping each domain-bin intersection were identified. The vessel area fraction per bin was defined as the total vessel area divided by the domain area within that bin. Vessel area fractions were smoothed using a rolling mean with a window size of 10 bins. The y-axis was scaled to [0, 1] to enable comparison across slides with different wall thicknesses.

### Age related vessel density changes

To test for age-associated changes in vessel subtype density, we fit a generalized linear model (GLM) with a Gamma distribution and log link function independently for each organ-vessel subtype combination. The model was specified as: density ∼ Age + Sex + log(Ischemic Time). Age was standardized (z-scored) for model fitting to improve numerical stability; the resulting coefficient was then rescaled to obtain the effect per year of age. The rate ratio per decade of aging was computed as exp(10 * beta), and the percent change per decade as (exp(10 * beta) - 1) * 100. Organ-vessel combinations with fewer than 10 observations were excluded. To account for multiple testing, p-values for the age coefficient were corrected within each organ using the Benjamini-Hochberg false discovery rate (FDR) procedure at a significance threshold of q < 0.05. This per-organ correction controls for the number of vessel subtypes tested within each organ rather than across all organ-vessel combinations jointly, reflecting the organ-stratified hypothesis structure.

A Gamma GLM with log link was fit to each organ-vessel combination, as described above, with the same model specifications. The model-predicted trend line was overlaid on the scatter plot, computed by evaluating the fitted model across the observed age range while holding sex at the modal category and log-transformed ischemic time at its mean (z = 0).

### Cell graph construction and topology analysis

To characterize tissue architecture at the whole-slide level, cell spatial proximity graphs were constructed for each WSI. Cell centroids from the largest tissue piece (identified via tissue segmentation with LazySlide^52^) were used as graph nodes. Edges were added between all pairs of cells whose centroids lay within 50 µm of each other, determined via KDTree query. Edge weights were recorded as Euclidean distances in both pixel and micrometer units.

From each cell graph, a set of topology features was extracted capturing global graph structure, node-level centrality distributions and spatial organization. Global features included node and edge counts, average degree, graph density, connected component statistics, largest connected component (LCC) size, Weisfeiler-Lehman color count, degree assortativity, k-core decomposition statistics (up to k = 10), edge counts (cliques_2) and triangle count (clique_3), the latter computed via the sparse identity (A·A)°A summed and divided by six for efficiency on large graphs. Community structure was assessed via Louvain clustering (resolution = 1.0) on the LCC, yielding modularity and number of communities. Node-level ranking features were summarized as distributional statistics (mean, standard deviation, minimum, maximum, 5th and 95th percentiles) for degree, PageRank, Katz centrality.

Spatial features included nearest-neighbor distance statistics (k = 1, 2, 3), mean effective radius, bounding box dimensions, spatial density and boundary statistics (distance to convex hull, boundary node fraction). Edge length distributions were similarly summarized. Features dependent on tissue size (node and edge counts, k-core bin counts, clique counts) were normalized by tissue area. Zero-variance and missing-value features were removed. For dimensionality reduction, features were scaled to zero mean and unit variance, followed by PCA. A shared nearest-neighbor graph was constructed using 10 neighbors and 20 principal components. To examine organ-specific structure, the same procedure was repeated independently within each organ.

### Cell community detection and quantification

To identify spatially coherent cellular neighborhoods within each tissue, community detection was performed on the cell graphs described above. The Leiden algorith^67^ was applied with resolution 1.5 and a fixed random seed (42), using a GPU-accelerated backend (cuGraph^68^) via NVIDIA H100 and L4 GPUs. Each resulting community represents a spatially contiguous group of cells with denser internal connectivity than expected by chance.

For each community, topological, compositional and spatial features were computed. Network features included community size, network density, degree assortativity, average clustering coefficient, mean degree and degree standard deviation, mean k-core number, boundary ratio and conductance. Cell type composition was characterized by the dominant cell type and its proportion, per-type percentages and spatial densities (cells per unit convex hull area), and diversity indices (Shannon entropy and Simpson index). Composition purity was defined as 1 minus the normalized Shannon entropy.

Spatial properties were derived from the convex hull of community node coordinates, including hull area, perimeter, isoperimetric quotient, PCA-based elongation, aspect ratio and compactness (mean distance to centroid normalized by the square root of hull area). A spatial coherence score (product of isoperimetric quotient and spatial density) and a combined unit quality score (weighted average of spatial coherence, composition purity and structural isolation) were computed for each community. Physical interaction distances within communities were summarized as the mean, standard deviation and median of edge lengths in micrometers.

### Functional tissue unit characterization and age-associated changes

To define organ-specific functional tissue unit (FTU) types, the community-level feature matrix was assembled into an AnnData object with community features as variables and per-community metadata (slide identity, organ, donor age, sex, ischemic time) as observations. Communities with fewer than 10 nodes were excluded. Spatial areas were converted from pixels to mm^2^ using the slide-level microns-per-pixel resolution. Muscle and connective tissue cell types were merged into a single connective category. For each organ independently, zero-sum features were removed, remaining features were scaled to zero mean and unit variance, and PCA was computed. A shared nearest-neighbor graph (10 neighbors, 20 PCs) was constructed, followed by UMAP embedding and Leiden clustering (igraph resolution = 0.8). PAGA trajectory analysis was performed on the resulting clusters. Each Leiden cluster within an organ was treated as a distinct FTU type.

To quantify age-associated changes in FTU abundance, the area fraction occupied by each FTU type per slide was computed as the summed convex hull area of all communities assigned to a given cluster, divided by the total tissue area of the slide. A generalized linear model (GLM) with Gamma family and log link was fit for each organ-FTU combination, with area fraction as the response and donor age as the primary predictor, controlling for sex and ischemic time. Area fractions were clipped to a minimum of 1e-6 to satisfy the Gamma distribution’s positive-support constraint. The percentage change per decade was computed as (exp(10 * beta) - 1) * 100. P-values for the age coefficient were corrected within each organ using the Benjamini-Hochberg FDR procedure at q < 0.05.

To assess age-associated changes in FTU features, each community feature was modeled independently within each organ-FTU combination using OLS regression with age as the predictor and sex and ischemic time as co-variates. Clusters with fewer than 20 communities were excluded. The percentage change per year was calculated as (beta / mean) * 100 and scaled to a per-decade estimate. P-values were corrected per cluster using Benjamini-Hochberg FDR.

### Compositional versus structural age-related changes in functional tissue units

To distinguish FTU abundance changes that can be explained by shifts in cell type composition from those reflecting structural tissue remodeling, we compared the observed FTU abundance change with the change expected from cell type density trends alone. For each FTU cluster within an organ, an expected compositional change was computed as the density-weighted sum of independently measured cell type density changes with age. Specifically, for each cell type present in a given FTU, its mean density within the FTU was multiplied by the corresponding age-associated percentage change per year (derived from organ-level OLS regression of cell type densities on age, as described above), and the products were summed and normalized by total cell density to yield a percentage expected change per year. The observed FTU abundance change (from the Gamma GLM described above) was then regressed against this expected compositional change using linear regression. Residuals from this regression represent FTU abundance changes not accounted for by cell type composition shifts. Per-organ residual summaries were computed as summed absolute, positive and negative residuals with standard errors. To identify cell types associated with unexplained FTU changes, FTUs were split into two groups based on the sign of their regression residual (expanding more than expected vs. declining more than expected). For each cell type, the mean absolute compositional contribution was computed within each group, and the difference between groups was used to identify cell types preferentially associated with unexplained expansion or decline.

### CONCH feature extraction and text-image similarity scoring

To characterize histological and pathological features of gastrointestinal tissues at the tile level, we employed the CONCH vision-language foundation model. Whole-slide images were tiled at 256 pixels using LazySlide, and CONCH tile-level visual embeddings were extracted. Three sets of curated text terms were used for text-image similarity scoring: pathology-related terms (raw cosine similarity, without softmax normalization), microanatomy domain terms (with softmax normalization for domain segmentation), and general histological terms (raw cosine similarity). Text embeddings were computed via the CONCH text encoder, and cosine similarity between tile image embeddings and text embeddings was calculated for each term. Per-slide features were obtained by averaging tile-level embeddings and similarity scores across all tiles.

### Differential cell type abundance in pathology versus healthy gastrointestinal tissue

To assess cell composition differences between pathology-annotated and healthy gastrointestinal slides, cell type densities (cells per mm^2^) were computed for stomach and esophageal mucosa samples using the MLP-predicted cell type labels. Cell densities were log-transformed (log1p), and differential abundance was tested using the Wilcoxon rank-sum test (scanpy’s rank_genes_groups) comparing pathology versus healthy slides, with Benjamini-Hochberg FDR correction. This analysis was performed per organ and across all organs combined.

### Domain-specific cell type analysis

To resolve pathology-associated changes at the level of tissue microanatomy, we leveraged domain annotations previously mapped onto GTEx gastrointestinal tissues^25^ (muscularis, submucosa, mucosa, lamina propria, blood vessels, hemorrhage, submucosal gland, gastric foveolae and gastric gland). Cells were assigned to domains based on spatial overlap between their centroids and the annotated domain polygons. Cell counts per domain and cell type were normalized by total slide area to yield domain-specific cell type densities (cells per mm ^2^). For stomach samples, gastric glands and gastric foveolae were aggregated into a virtual “mucosa” domain to enable cross-organ comparison with esophageal mucosa. For each domain-cell type combination, the log2 fold change between pathology and healthy groups was computed (with a pseudo count of 1e-6), and significance was assessed using the Mann-Whitney U test with BH-FDR correction. The immune-to-epithelial ratio was computed per domain as log2((immune density + 1e-6) / (epithelial density + 1e-6)). A mean effect size heatmap across shared domains was derived by averaging per-organ log2 fold changes and taking the maximum (most conservative) adjusted p-value across organs.

### Vision-language characterization of pathological changes

To identify histological features associated with pathology in an unsupervised manner, mean CONCH tile embeddings per slide were used for PCA, neighbor graph construction and UMAP visualization using scanpy ^54^ with default parameters. For text-image similarity analysis, histological structure terms (mucosa, submucosa, muscle, etc.) were excluded from the general terms set to focus on pathological features. For each text term, the mean similarity difference between pathology and healthy groups was computed, and significance was assessed using the Wilcoxon rank-sum test with per-tissue BH-FDR correction.

## Data availability

GTEx whole slide images are available from its portal (https://gtexportal.org) With donor-specific demographic and clinical information available from dbGaP at http://www.ncbi.nlm.nih.gov/gap through dbGaP accession number phs000424.v9.p2.c1. Predicted cell phenotype identities, spatial coordinates, cell type densities, and tissue graphs from the histomorphological atlas are available at https://huggingface.co/datasets/RendeiroLab/histomorphological-cell-atlas.

## Code availability

The source code is provided in the Supplementary Information files for reviewers and will be publicly available after peer review at the GitHub repository: https://github.com/rendeirolab/CINage.

## Acknowledgments

The Rendeiro group was supported by Angelini Ventures S.p.A. Rome, Italy and funding from the European Research Council (ERC) under the European Union’s Horizon Europe research and innovation programme (grant agreement no. 101220825). The Genotype-Tissue Expression (GTEx) Project was supported by the Common Fund of the Office of the Director of the National Institutes of Health, and by NCI, NHGRI, NHLBI, NIDA, NIMH, and NINDS. The datasets used for the analyses described in this manuscript were obtained from dbGaP at http://www.ncbi.nlm.nih.gov/gap through dbGaP accession number phs000424.v9.p2.c1. We thank the IT team at CeMM for access and maintenance of the CeMM HPC cluster, and Barbara Maier for feedback on the manuscript.

