## Supplementary Figures for "A single-cell view of human tissue aging reveals architectural decline beyond cellular composition"

**Figure S1, related to Figure 1: Dataset overview. a)** Overview of sample distribution of the whole dataset, left: number of slides per tissue; middle: number of slides per tissue and age bracket; top-right: sex distribution; bot-right: age and sex distribution. **b)** Overview of the dataset balanced for age, sex and organ, used for cell type phenotyping and subsequent quantification across age, same as in a).

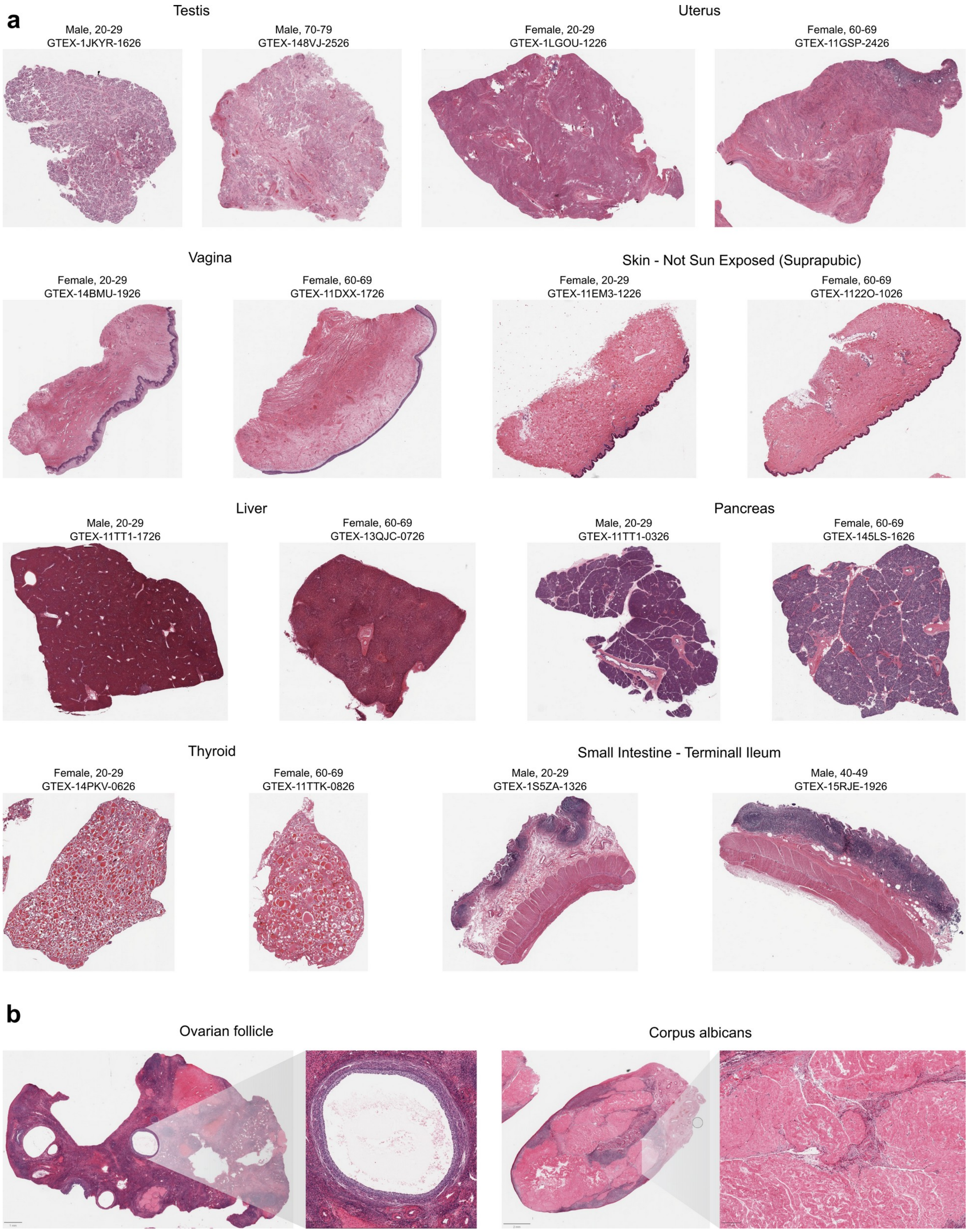


**Figure S2, related to Figure 1: Histological signatures of age driven changes**. **a)** Examples of histopathology scans for different organs and age brackets. **b)** Zoomed in examples of microanatomical domains in the ovary, left: ovarian follicle; right: corpus albicans.

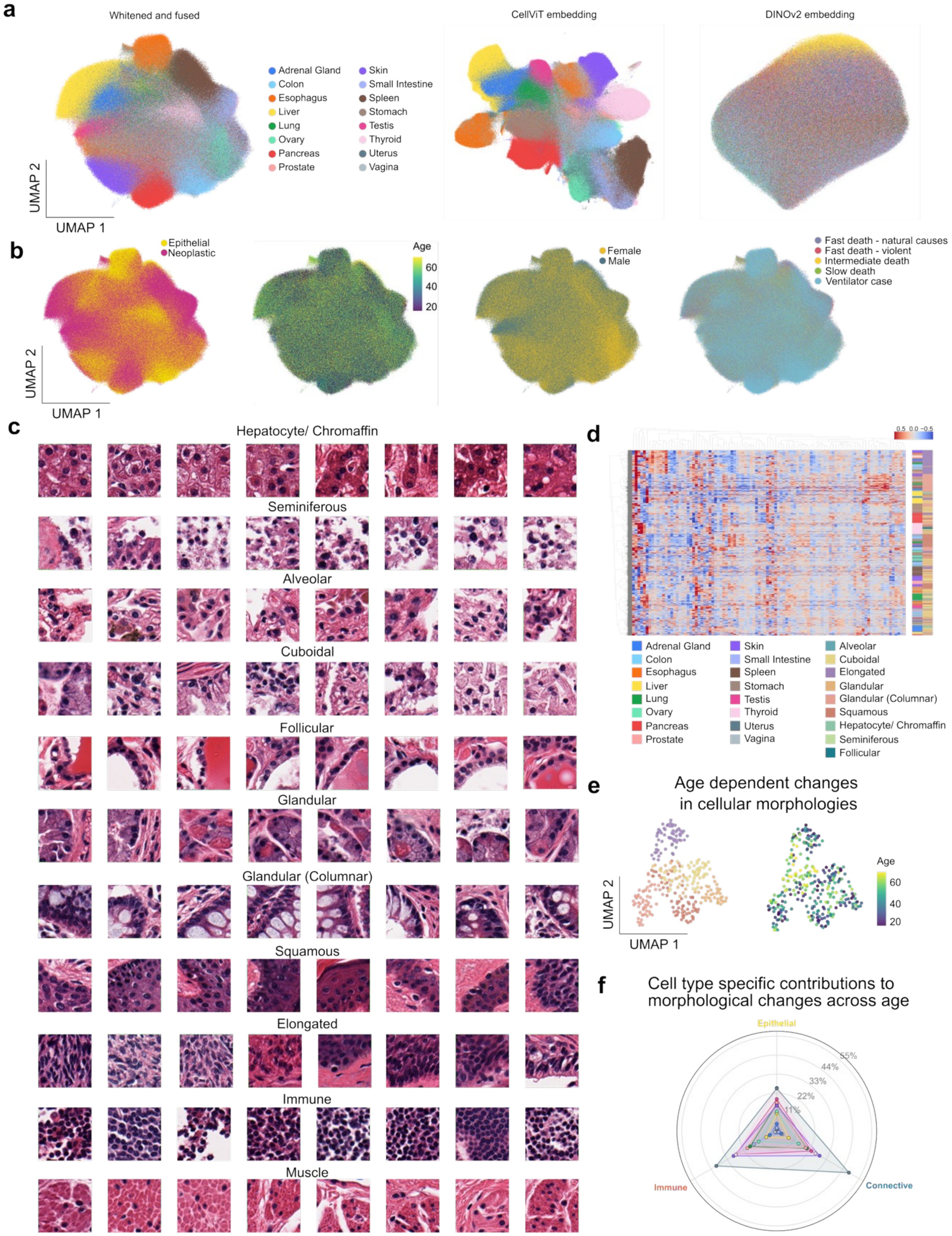


**Figure S3, related to Figure 2: Epithelial subtype phenotyping with multi-modal features. a)** UMAP embedding of features from combined (fused and whitened) or independent CellViT and DINOv2 cells, classified as epithelial and neoplastic. **b)** UMAP embedding of fused and whitened features colored by different variables. **c)** Exemplary cell images, centered around the classified cells, for all the identified phenotypes. **d)** Clustermap of mean slide and cell type aggregated fused and whitened features. **e)** UMAP of slide-level mean epithelial subtype features in the vagina (immune and muscle cells excluded). **f)** Polar plot of the mean percentage of CellViT and DINOv2 embedding dimensions significantly changing with age (FDR < 0.05) per organ and cell compartment (Epithelial, Connective, Immune). Each triangle represents one organ; radial extent indicates mean % features age-associated, averaged across cell types within each compartment. Organ colors as in d).

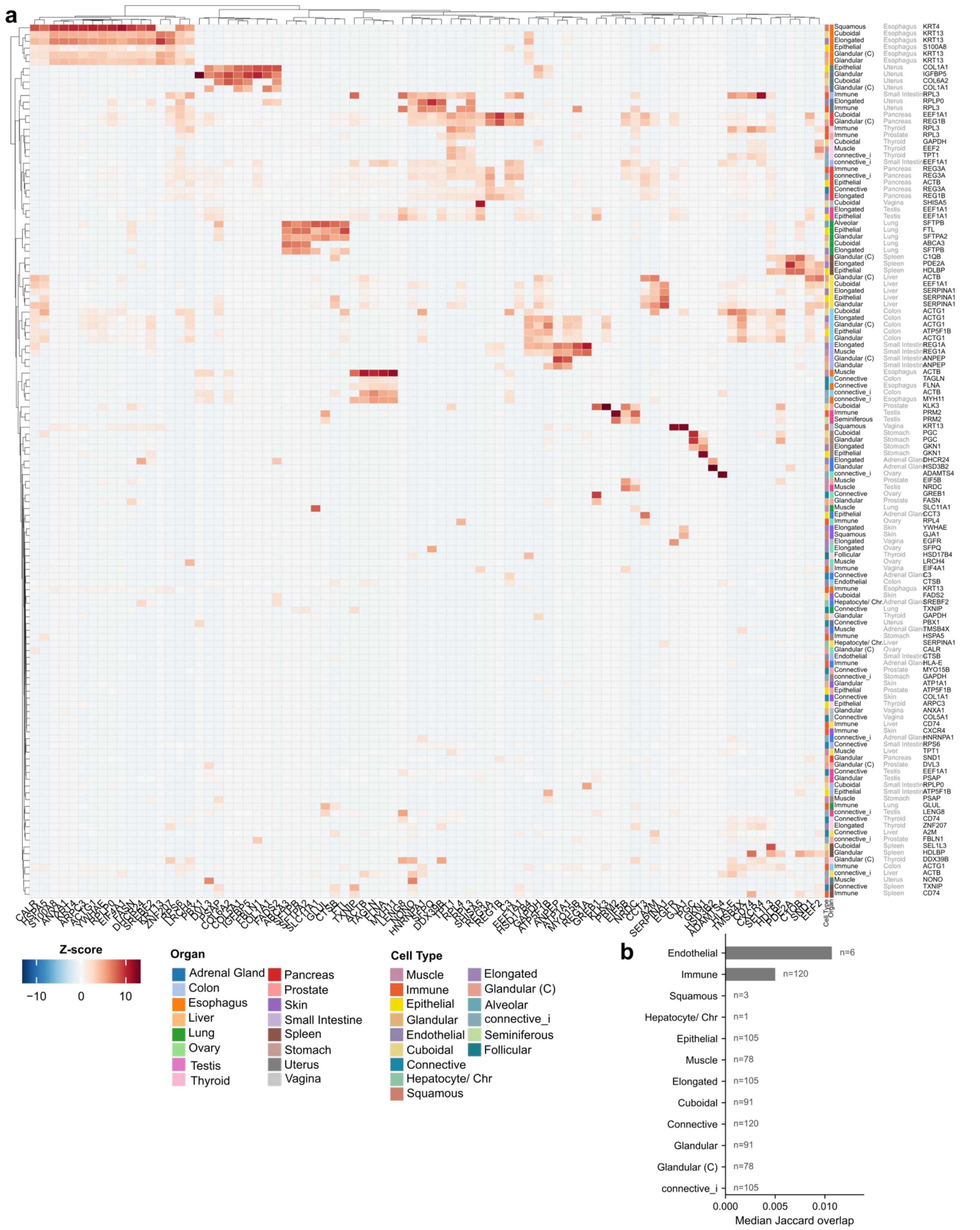


**Figure S4, related to Figure 3: Association of molecular profiles for histological phenotypes. a)** Heatmap with the union of top associated genes per cell type and organ, showing the specificity of molecular profiles. **b)** Jaccard index across organs for each morphological profile, showing that inferred molecular profiles are highly specific within each cell type across for all but two cell types: endothelial and immune cells.

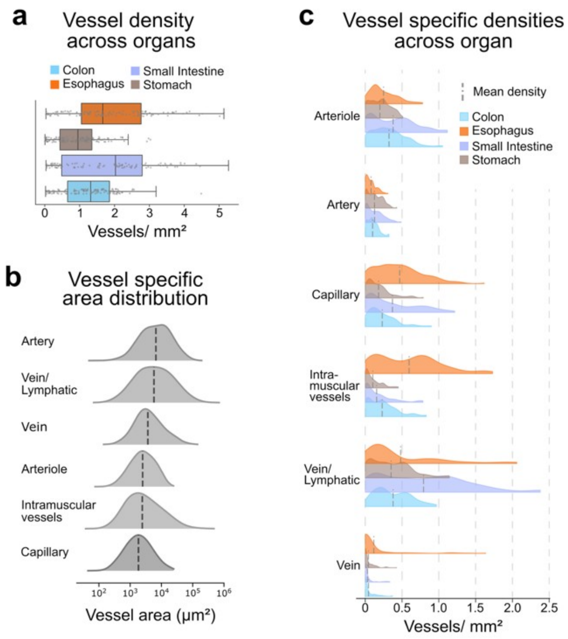


**Figure S5, related to Figure 4: Gastrointestinal vessel area, composition, density, and CONCH text-image similarity validation.** **a)** Total vessel density per organ (vessels per mm2). **b)** Ridgeline plots of vessel subtype densities across organs. **c)** Vessel polygon area distributions by vessel subtype (log-scaled). **d)** Heatmap of mean CONCH text-image similarity scores between vessel subtypes and curated histological text descriptors (phenotype-specific terms and background tissue terms as negative controls).

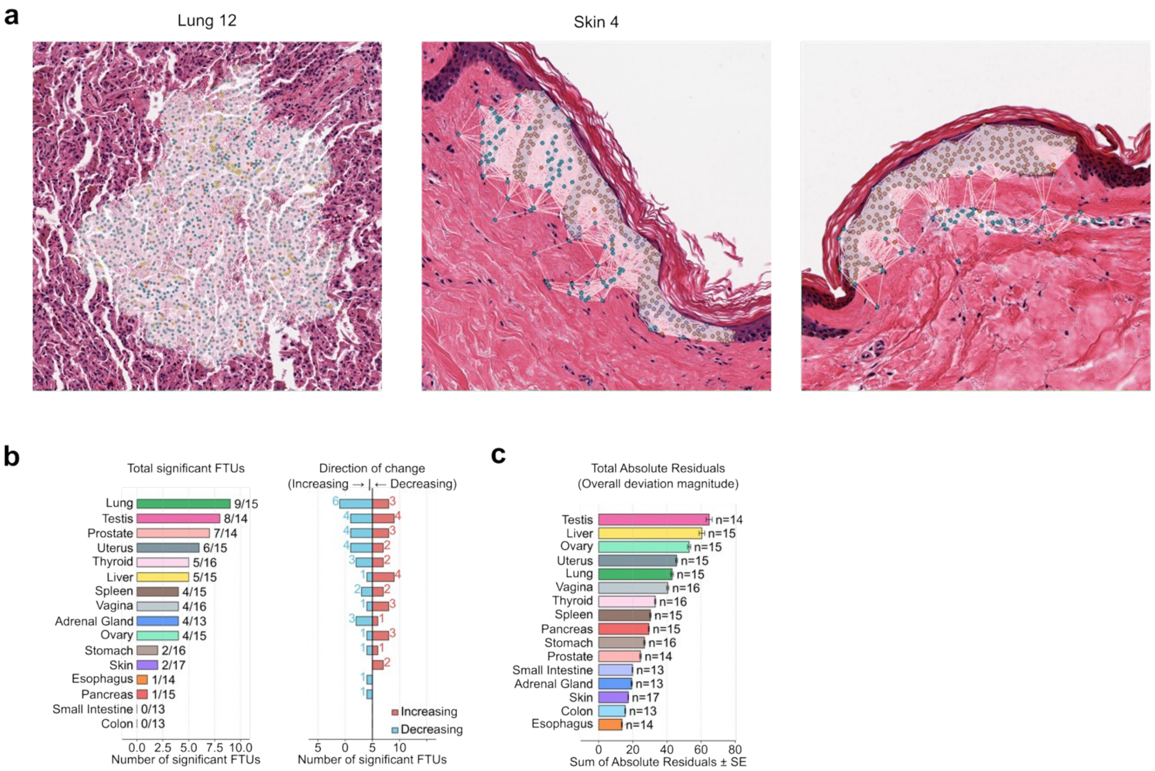


**Figure S6, related to Figure 5: Age dependent changes in FTU structure. a)** Exemplary FTUs of the lung and skin, strongly affected by age-related changes. **b)** Per-organ summary of significant FTU abundance changes with age and directional changes. **c)** Per-organ summary of total absulte residuals, where FTU abundance changes exceed compositional expectations.

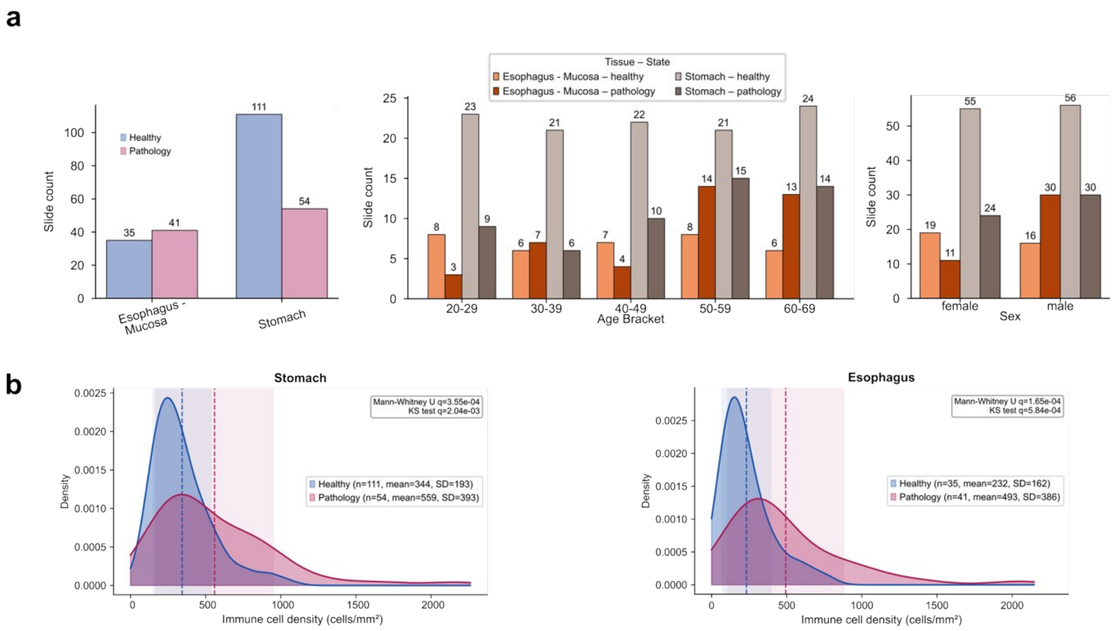


**Figure S7, related to Figure 6: Gastrointestinal pathology cohort overview and per-organ immune density distributions. a)** Sample distribution of the gastrointestinal pathology cohort across organs and pathology status. **b)** Kernel density estimates of immune cell density in healthy versus pathology slides, shown per organ (related to Fig. 5f). Dashed lines = group means; shaded bands = ± 1 SD. p-values: Mann-Whitney U, BH-corrected.
